# The chitin receptor-interacting protein LIK1 regulates extracellular ATP signaling via interaction with P2K1 in *Arabidopsis thaliana*

**DOI:** 10.64898/2026.04.08.716789

**Authors:** Jinrong Wan, Mengran Yang, Jae Song, Chunhui Xu, Sung-Hwan Cho, Mowei Zhou, Ljiljana Pasa-Tolic, Bing Yang, Dong Xu, Gary Stacey

## Abstract

Previously, the chitin receptor-interacting protein kinase LIK1 (LysM receptor kinase 1/CERK1-interacting kinase) was shown to play an important role in regulating chitin signaling and plant defense. A limited proteolysis proteomics study revealed several LIK1-derived peptides that showed differential abundance between ATP-treated and mock-treated Arabidopsis samples, suggesting a possible involvement of LIK1 in extracellular ATP (eATP) signaling. To explore this possibility, *LIK1* mutants were obtained and examined for their response to ATP. The results showed that mutations in *LIK1* significantly reduced the expression of eATP-responsive genes. In addition, LIK1 was found to interact with the eATP receptor P2K1 and to be phosphorylated by it. The LIK1 protein was localized to the plasma membrane and its gene expression appeared to be ubiquitous. Collectively, these findings indicate that LIK1 not only contributes to chitin signaling but also participates in eATP signaling, highlighting its potential role as a shared component in multiple signaling pathways to regulate plant responses to diverse internal and external cues.

## 1. Introduction

Plants are continually exposed to a wide range of biotic and abiotic challenges and have evolved complex, interconnected signaling pathways to perceive and respond to these environmental stimuli. One well-characterized defense mechanism is the chitin-mediated signal transduction pathway, which enables plants to detect fungal pathogens by recognizing chitin, a major component of fungal cell walls, through the LysM receptor kinase LysM RLK1 (also known as CERK1) (Miya et al., 2007; Wan et al., 2008). Some key components of this pathway have been identified, including LIK1 (LysM RLK1-interacting protein kinase 1), a protein discovered in a previous mutant screen (Le et al., 2014).

LIK1 is a leucine-rich repeat (LRR)-like kinase predicted to contain a signal peptide, an extracellular LRR domain, a transmembrane domain, and an intracellular kinase domain. Mutations in *LIK1* enhanced the expression of chitin-responsive genes upon chitin treatment while suppressing the expression of genes involved in the jasmonic acid (JA) signaling pathway. Consequently, *LIK1* mutant plants exhibited increased resistance to the hemibiotrophic bacterial pathogen *Pseudomonas syringae* but heightened susceptibility to the necrotrophic fungal pathogen *Sclerotinia sclerotiorum* compared to wild-type plants (Le et al., 2014). Our earlier work also showed that LIK1 interacted with and was phosphorylated by LysM RLK1/CERK1 (Le et al., 2014). These findings suggest that LIK1 plays an important regulatory role in plant defense by modulating both chitin and JA signaling pathways.

Beyond the findings reported by Le et al. (2014), little was previously known about the function of LIK1. However, a recent limited proteolysis (LiP) proteomics study, conducted to identify additional components involved in extracellular ATP (eATP) signaling, found that several LIK1-derived peptides showed differential abundance between ATP-treated and mock-treated Arabidopsis samples (Table S1). In addition, a recent study demonstrated that LIK1 co-precipitated with the eATP receptor P2K1 in Arabidopsis (Sowders et al., 2024). Together, these findings suggest that LIK1 may also play a role in eATP signaling.

eATP is ATP released by plant cells into the extracellular space in response to internal or external stimuli (Tanaka et al., 2014). It functions as a key signaling molecule involved in various aspects of plant growth, development, and stress responses (Choi et al., 2014a, b; Yuan et al., 2025). P2K1, a lectin receptor-like kinase, was the first eATP receptor identified in plants and plays a central role in perceiving and mediating eATP-triggered signal transduction (Choi et al., 2014a). Studies have shown that eATP can act as a damage-associated molecular pattern (DAMP) to trigger defense responses to cell wounding (Choi et al., 2014b; Myers et al., 2022; Tanaka et al., 2014; Tanaka and Heil, 2021). Numerous studies also demonstrated that JA is involved in plant responses to injury or wounding caused by herbivores, pathogens, and mechanical damage (Kimberlin et al., 2022; Koo and Howe, 2009; Ma et al., 2025). The gene expression profiles induced by JA and ATP significantly overlap (Choi et al., 2014a). Furthermore, a recent study revealed that JA can prime plant responses to eATP via P2K1 (Jewell et al., 2024). Therefore, the signaling pathways mediated by eATP and JA are likely interconnected.

Given that LIK1 positively regulated the JA pathway (Le et al., 2014), exhibited differential responses to ATP treatment in our LiP proteomics study compared to the mock (buffer) control, and co-precipitated with the eATP receptor P2K1 (Sowders et al., 2024), we hypothesized that LIK1 may also participate in eATP signaling. To explore this possibility, we obtained *LIK1* mutant lines and examined their responses to ATP. Loss-of-function mutations in *LIK1* significantly reduced the expression of eATP-responsive genes. We further demonstrated that LIK1 was localized to the plasma membrane and interacted with and was phosphorylated by P2K1. Collectively, these findings support a role for LIK1 in eATP signaling, in addition to its function in chitin signaling, suggesting that LIK1 may act as a multifunctional signaling component that integrates diverse internal and external signals to regulate plant growth, development, and stress responses.

## 2. Materials and methods

### 2.1. Plant materials, growth conditions, and treatment

The *Arabidopsis thaliana* T-DNA insertion mutant SALK_030855 (named *lik1-1* in this study) was obtained from the Arabidopsis Biological Resource Center (ABRC) and genotyped using primers listed in Table S2.

*lik1-2* and *lik1-3* mutants were generated using the CRISPR-Cas9 technology. The primers used for this purpose are listed in Table S2.

For gene expression analysis, seeds were surface sterilized and sown on half-strength Murashige and Skoog (MS) medium supplemented with 1% (w/v) sucrose and 0.5% (w/v) phytagel, adjusted to pH 5.7. Plates were stratified at 4□°C for 3 days in darkness and then transferred to a growth chamber to grow vertically under long-day conditions (16 h light/8 h dark cycle) at 22□°C, with a light intensity of 100 μE□cm−^2^□s−^1^. Seven-day-old seedlings were treated with 200 µM ATP for 30 minutes. As a negative control, seedlings were treated with an equivalent volume of 50 mM MES buffer (pH 5.7, used for ATP solution preparation). After treatment, whole seedlings were harvested and immediately frozen in liquid nitrogen for subsequent RNA isolation.

### 2.2. RNA isolation, cDNA synthesis, RT-PCR, and quantitative RT-PCR

Total RNA was extracted from whole *Arabidopsis* seedlings using the RNeasy Plant Mini Kit (QIAGEN, Hilden, Germany), following the manufacturer’s instructions. First-strand complementary DNA (cDNA) was synthesized using the RNA to cDNA EcoDry™ Premix with Random Hexamers (TaKaRa Bio Inc., Kusatsu, Shiga, Japan), according to the manufacturer’s protocol.

To assess *LIK1* transcript levels in both the *lik1-1* mutant and wild-type plants, RT-PCR was performed using gene-specific primers listed in Table S2. The *SAND* gene (*At2g28390*) was used as an endogenous control. RT-PCR was carried out using PrimeSTAR GXL DNA Polymerase (TaKaRa Bio Inc., Kusatsu, Shiga, Japan) under the following cycling conditions: 30 cycles of 98°C for 10 sec, 55°C for 15 sec, and 68°C for 1 min, followed by a final extension at 68°C for 3 min. The resulting PCR products were resolved on a 1% agarose gel containing ethidium bromide and visualized under UV light for comparison.

For quantitative RT-PCR (qRT-PCR), transcript levels of *WRKY53* (*At4g23810*), *CPK28* (*At5g66210*), *ATPR2* (*At3G44870*), *CML39* (At3g22910), and *SAND* (*At2g28390*) were quantified using a 7500 Real-Time PCR System (Applied Biosystems, Waltham, MA) and SYBR Green Master Mix (Applied Biosystems, Waltham, MA). Primer sequences used were listed in Table S2. The relative fold change of the target gene, normalized to the expression of the endogenous control gene *SAND* due to its highly stable expression (Czechowski et al. 2005) and relative to its gene expression in the mock treated control sample, was calculated as described (Livak and Schmittgen, 2001).

### 2.3. Split-luciferase complementation assay

The full-length coding sequence (CDS, without the stop codon) of *LIK1* was cloned into the entry vector pDNOR/Zeo and then into the vectors pCambia-NLuc (abbreviated as NLuc) and pCambia-CLuc (abbreviated as CLuc) (Chen et al., 2008) using Gateway cloning strategies (Curtis and Grossniklaus, 2003) to generate the LIK1-NLuc and LIK1-CLuc constructs, respectively. The P2K1-CLuc and P2K1-NLuc constructs were obtained from a previous publication (Pham et al., 2020). These constructs were electroporated into *Agrobacterium tumefaciens* strain GV3101.

*Tobacco (Nicotiana benthamiana)* leaves were co-infiltrated with *Agrobacterium* strain pairs harboring LIK1-NLuc/P2K1-CLuc, LIK1-NLuc/CLuc (Negataive control), NLuc/P2K1-CLuc (negative control), or NLuc/CLuc (empty vector control), together with *Agrobacterium* strain expressing the RNA silencing suppressor p19 (Silhavy et al., 2002), as described previously (Li, 2011). Three days post-infiltration, the leaves were collected and sprayed with 1 mM luciferin (Gold Biotechnology, Inc., St. Louis, MO, USA) and bioluminescence was detected using a CCD imaging system (Photek 216).

### 2.4. Bimolecular fluorescence complementation (BiFC) assay

To assess protein–protein interactions between LIK1 and P2K1, bimolecular fluorescence complementation (BiFC) assays were performed. The full-length coding sequence (CDS) of *LIK1* (excluding the stop codon) was cloned into pAM-PAT-35S:YFPn:GW (abbreviated as NYFP) and pAM-PAT-35S:YFPc:GW (abbreviated as CYFP) vectors to generate LIK1-NYFP and LIK1-CYFP, respectively, using Gateway cloning strategies (Curtis and Grossniklaus, 2003).

The full-length CDS of *P2K1* (without the stop codon) had previously been cloned into the same vectors to generate P2K1-NYFP and P2K1-CYFP (Chen et al., 2017; Cho et al., 2022).

All constructs were electroporated into *Agrobacterium tumefaciens* strain GV3101. Transformed strains were cultured for one day, pelleted, and resuspended in 10□mM MgCl_2_. After pretreatment with 40□µM acetosyringone for 2 hours at room temperature, bacterial cultures were mixed in equal proportions with *Agrobacterium* expressing the RNA silencing suppressor p19 (Silhavy et al., 2002), to a final OD_600_ of 0.6.

The following combinations were infiltrated into 4-week-old *Nicotiana benthamiana* leaves using a needleless syringe: LIK1-NYFP/LIK1-CYFP, LIK1-NYFP/P2K1-CYFP, P2K1-NYFP/LIK1-CYFP, LIK1-NYFP/CYFP (negative control), NYFP/LIK1-CYFP (negative control), P2K1-NYFP/CYFP (negative control), NYFP/P2K1-CYFP (negative control), and NYFP/CYFP ((empty vector control).

Three days after infiltration, the leaf sections were excised and analyzed using confocal microscopy to detect YFP fluorescence to infer protein–protein interaction.

### 2.5. Protein expression

The coding sequence (CDS) of the LIK1 kinase domain was cloned into the pRSFDuet-1_VNp-His6 vector (Eastwood et al., 2023) for protein expression. A kinase-dead mutant version, LIK1KD-D798A, was also generated by site-directed mutagenesis (Le et al., 2014) and cloned into the same vector at the BglII and NheI restriction sites. The primers used for cloning were listed in Table S1. The confirmed constructs were transformed into *ArcticExpress* (DE3) *E. coli* strain and protein expression was performed according to the manufacturer’s protocol (Agilent Technologies, Inc., Santa Clara, CA). Recombinant proteins were purified using TALON Metal Affinity Resin (Clontech Laboratories, Inc., a Takara Bio Company, Mountain View, California, USA) following the manufacturer’s instructions.

For the GST-tagged constructs, the kinase domain of P2K1 (GST-P2K1-KD) and its kinase-dead variant GST-P2K1-D572N-KD (d1-1) were fused to GST in the pGEX-5X-1 vector (GE Healthcare), as described previously (Choi et al., 2014a). The constructs were transformed into *Rosetta* (DE3) *E. coli* strain, which expresses the YopH tyrosine phosphatase to aid kinase protein expression. GST-tagged proteins were purified using Glutathione Resin (GenScript, Nanjing, China) following the manufacturer’s protocol.

### 2.6. In vitro kinase assay

The purified recombinant proteins were subjected to an in vitro kinase assay following the protocol described by Cho et al. (2022). Briefly, kinase reactions were performed in 50 mM Tris-HCl, pH 7.5, 10 mM MgCl2, 10 mM MnCl_2_, 5 mM EGTA, 100 mM NaCl, 1 mM DTT with both radiolabeled and unlabeled ATP, and incubated for 1.5 hours at 30 °C. The reaction products were separated by electrophoresis on a 12.5% SDS-polyacrylamide gel. Phosphorylated proteins were then detected by autoradiography (Typhoon FLA9000, GE Healthcare).

### 2.7. Subcellular localization of LIK1

To determine the subcellular localization of LIK1, the coding sequence of *LIK1* (excluding the stop codon) was cloned into the binary vector pMDC83 to generate a LIK1-GFP fusion construct via Gateway cloning (Curtis and Grossniklaus, 2003). The LIK1-GFP construct and a plasma membrane marker (PM-mCherry) were electroporated into *Agrobacterium tumefaciens* strain GV3101, respectively. The transformed *Agrobacterium* cultures were grown overnight, pelleted, and resuspended in 10□mM MgCl_2_. Following a 2-hour pretreatment with 40□µM acetosyringone at room temperature, the LIK1-GFP and PM-mCherry strains were mixed in equal proportions with *Agrobacterium* C58C1 expressing the silencing suppressor HC-Pro (Llave et al., 2000), to a final optical density of OD_600_ = 0.6. The resulting bacterial suspension was infiltrated into leaves of 4-week-old *Nicotiana benthamiana* plants using a needleless syringe. Three days post-infiltration, leaf sections were collected and observed under a confocal microscope to examine the localization of LIK1-GFP relative to the PM-mCherry marker.

For plasmolysis assays, infiltrated leaves were treated with 3□M NaCl for approximately 15 minutes before imaging. Retraction of the LIK1-GFP signal from the cell wall would be used to confirm its localization to the plasma membrane.

### 2.8. GUS Staining

To analyze the expression pattern of *LIK1*, a 2470-bp sequence upstream of the *LIK1* start codon was amplified from the genomic DNA of *Arabidopsis thaliana* Col-0 plants and cloned into the binary vector pMDC162 to generate the *LIK1* promoter-GUS fusion construct, using Gateway cloning strategies (Curtis and Grossniklaus, 2003). The resulting construct was electroporated into *Agrobacterium tumefaciens* strain AGL1 and used to transform *Arabidopsis* Col-0 plants via the floral dip method (Clough and Bent, 1998). Transgenic lines were selected on medium containing hygromycin.

Histochemical GUS staining was performed as previously described (Jefferson et al., 1987) to assess promoter activity.

### 2.9. Limited Proteolysis Proteomics

Ten-day-old seedlings overexpressing P2K1 were treated with ATP at 200 µM or buffer (as the negative control) for 15 minutes. The total membrane proteins were isolated from the treated seedlings according to the procedure described by Zhou et al. (2020). The total membrane proteins were then processed for mass spectrometry analysis following the steps described by Cappelletti et al. (2021).

## 3. Results

### 3.1. Mutations in LIK1 reduce the expression of eATP-responsive genes following ATP treatment

To investigate whether LIK1 is involved in eATP signaling, we first obtained a T-DNA insertion mutant SALK_030855 (named *lik1-1* hereafter, Fig. S1) and assessed its response to ATP treatment. As shown in Fig. S2, the mutation in *LIK1* significantly reduced the expression of the previously identified ATP-responsive genes *CPK28* and *WRKY40* (Choi et al., 2014a) compared to wild-type plants. To validate this result, we obtained additional *lik1* mutant lines *lik1-2* and *-3* using the CRISPR-Cas9 technology (Figs. S3-6). Consistent with the T-DNA mutant phenotype, these CRISPR-Cas9-derived lines also showed reduced the expression of *CPK28* and *WRKY40* as weak as genes *ATPR2* and *CML39* (Sowders et al., 2024) upon ATP treatment (Fig. 1). These findings support a possible role for LIK1 in regulating eATP signaling in addition to its established role in chitin signaling.

**Fig. 1.**
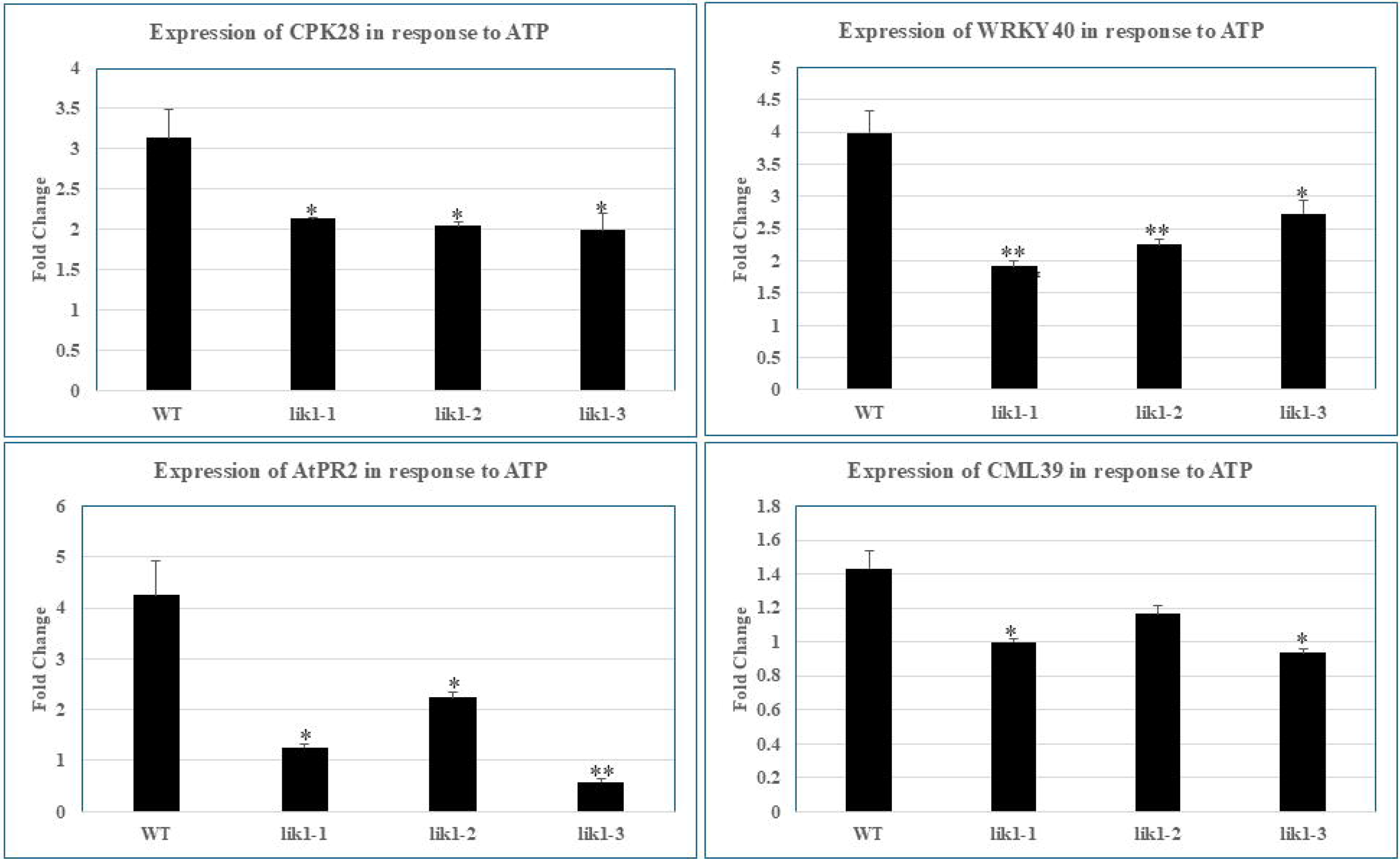
qRT-PCR analysis of ATP-responsive genes in the *LIK1* mutants. The genes examined were CPK28, WRKY40, ATPR2, and CML39. One-week-old seedlings were treated with 200 µM ATP or mock control (MES buffer) for 30 minutes. The relative expression (fold change) of a target gene was obtained by comparing the ATP-treated samples to mock controls, following normalization to the reference gene *SAND*. Data represented the mean ± SE (standard error) of four biological replicates. Statistical significance (between the WT and the mutants) was determined using a Student’s t-test:*, *P* value <0.05; **, *P* value <0.01.

### 3.2. LIK1 self-associates and interacts with P2K1 at the plasma membrane

Given that LIK1 is predicted to localize to the plasma membrane (PM) (Hooper et al., 2014 & 2017) and P2K1 is a membrane-bound eATP receptor, we first investigated whether these two proteins interact. As shown in Fig. 2, a split-luciferase complementation assay performed in *Nicotiana benthamiana* leaves indicated that LIK1 and P2K1 interacted, although the signal suggested a relatively weak interaction. This interaction was further supported by bimolecular fluorescence complementation (BiFC) assays, which also revealed a weak but consistent interaction between LIK1 and P2K1 (Fig. 3 and Fig. S7). Interestingly, LIK1 also exhibited strong self-association in the BiFC assay (Fig. 4), suggesting it can also form homodimers, in addition to forming a complex with P2K1 as well as LysM RLK1/CERK1. As noted in the Introduction, the finding that LIK1 co-precipitated with P2K1 in *Arabidopsis* (Sowders et al., 2024) and our finding that LIK1 was phosphorylated by P2K1 (see below) further supports the interaction between these two proteins.

**Fig. 2.**
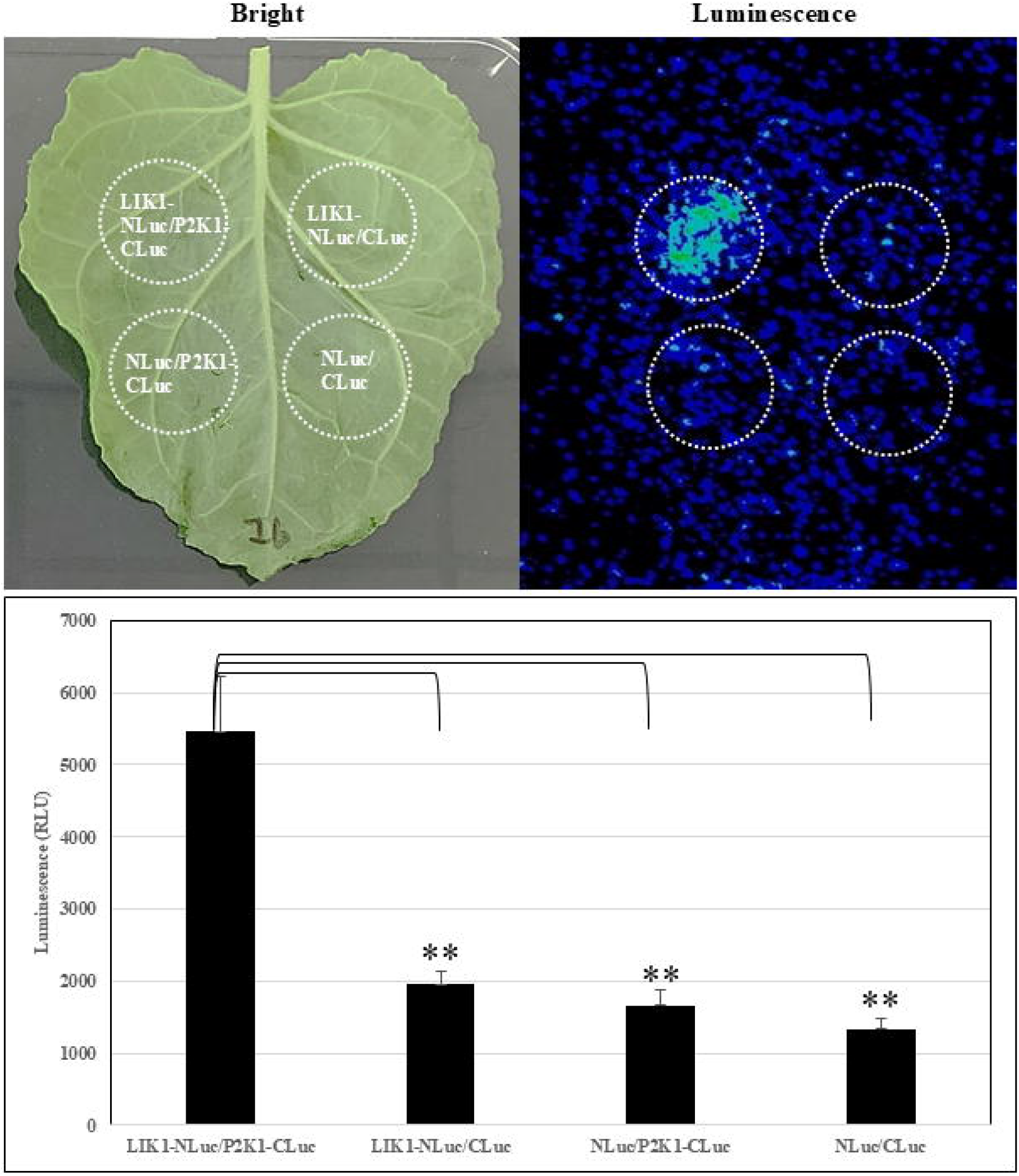
Split-luciferase assay showing the LIK1 and P2K1 interaction in tobacco leaves. Tobacco (*Nicotiana benthamiana)* leaves were co-infiltrated with the following agrobacterial strain pairs containing: LIK1-NLuc/P2K1-CLuc, LIK1-Nluc/Cluc, Nluc/P2K1-CLuc, or Nluc/CLuc, as indicated in the figure. The Circles indicate the leaf areas that were infiltrated with these *Agrobacterium* stains. (A) The bright and luminescence images of the infiltrated leaf. (B) Quantification of the luminescence from each infiltrated area. Asterisks indicate a statistically significant difference (*n* = 6 leaves; T-test: **, *P* < 0.01).

**Fig. 3.**
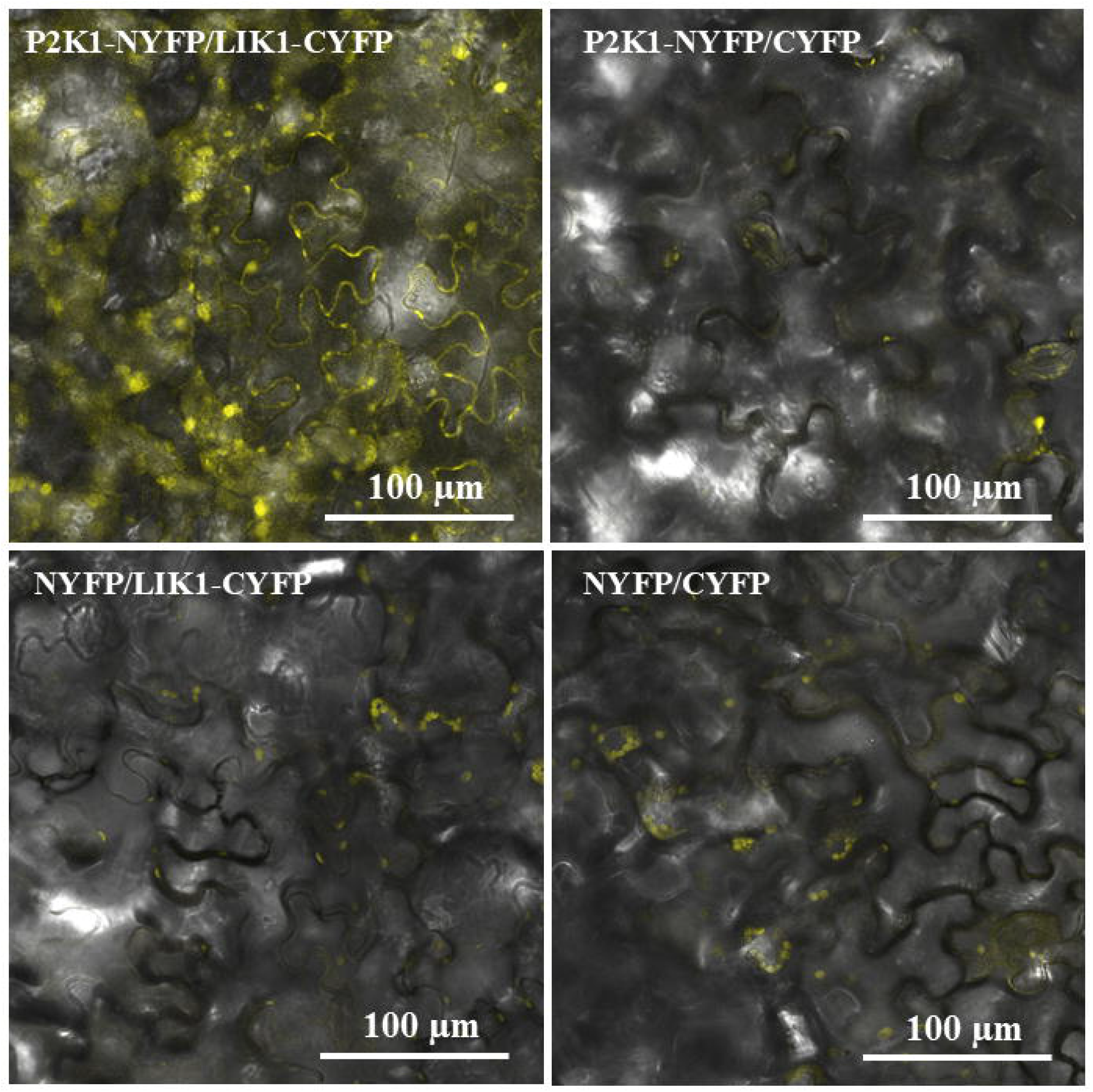
P2K1-LIK1 interaction in the bimolecular fluorescence complementation (BiFC) assay. Tobacco (*Nicotiana benthamiana)* leaves were co-infiltrated with the following agrobacterial strain pairs containing: P2K1-NYFP/LIK1-CYFP, P2K1-NYFP/CYFP, NYFP/LIK1-CYFP, NYFP/CYFP, as indicated in the figure. The infiltrated leaves were observed ∼ three days later under a confocal microscope. Scale bars = 100 μm.

**Fig. 4.**
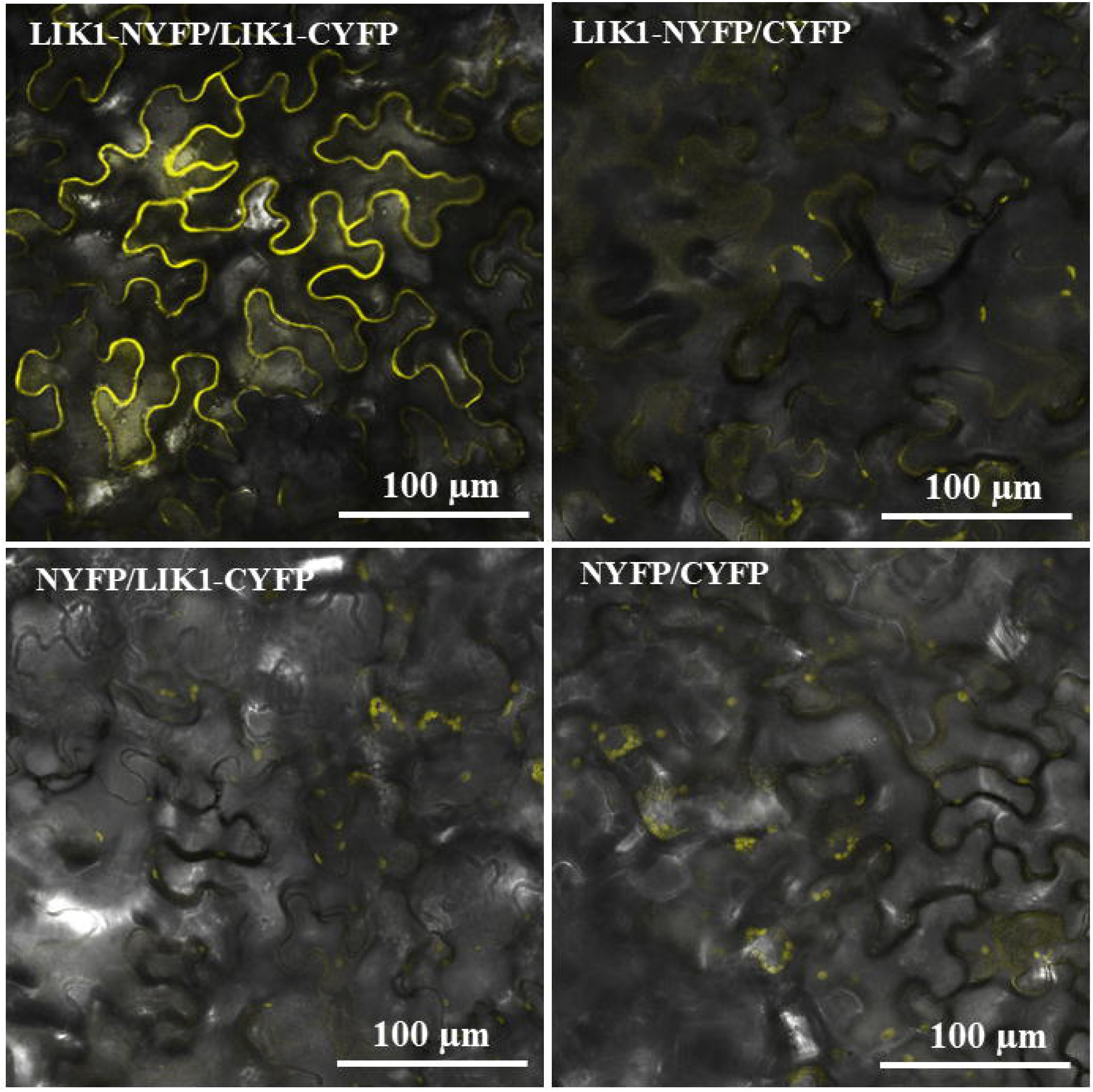
LIK1-LIK1 interaction in the bimolecular fluorescence complementation (BiFC) assay. Tobacco (*Nicotiana benthamiana)* leaves were co-infiltrated with the following agrobacterial strain pairs containing: LIK1-NYFP/LIK1-CYFP, LIK1-NYFP/CYFP, NYFP/LIK1-CYFP, and NYFP/CYFP, as indicated in the figure. The infiltrated leaves were observed ∼ three days later under a confocal microscope. Scale bars = 100 μm.

### 3.3. LIK1 is phosphorylated by P2K1

Given that P2K1 is capable of phosphorylating various substrate proteins, as demonstrated in our previous studies (Jorge et al., 2024; Kim et al., 2025) and P2K1 interacts with LIK1, we performed *in vitro* kinase assays using purified recombinant kinase domains of both wild-type and kinase-dead P2K1, along with wild-type and mutant LIK1 kinase domains. As shown in Fig. 5, the wild-type P2K1 kinase domain was able to phosphorylate both the wild-type and mutant LIK1 kinase domains. In repeated assays, no autophosphorylation of LIK1KD was observed, suggesting that LIK1 either lacks intrinsic kinase activity or has very low activity under our current assay conditions. These findings indicate that LIK1 not only interacted with P2K1 but was also phosphorylated by this receptor kinase protein.

**Fig. 5.**
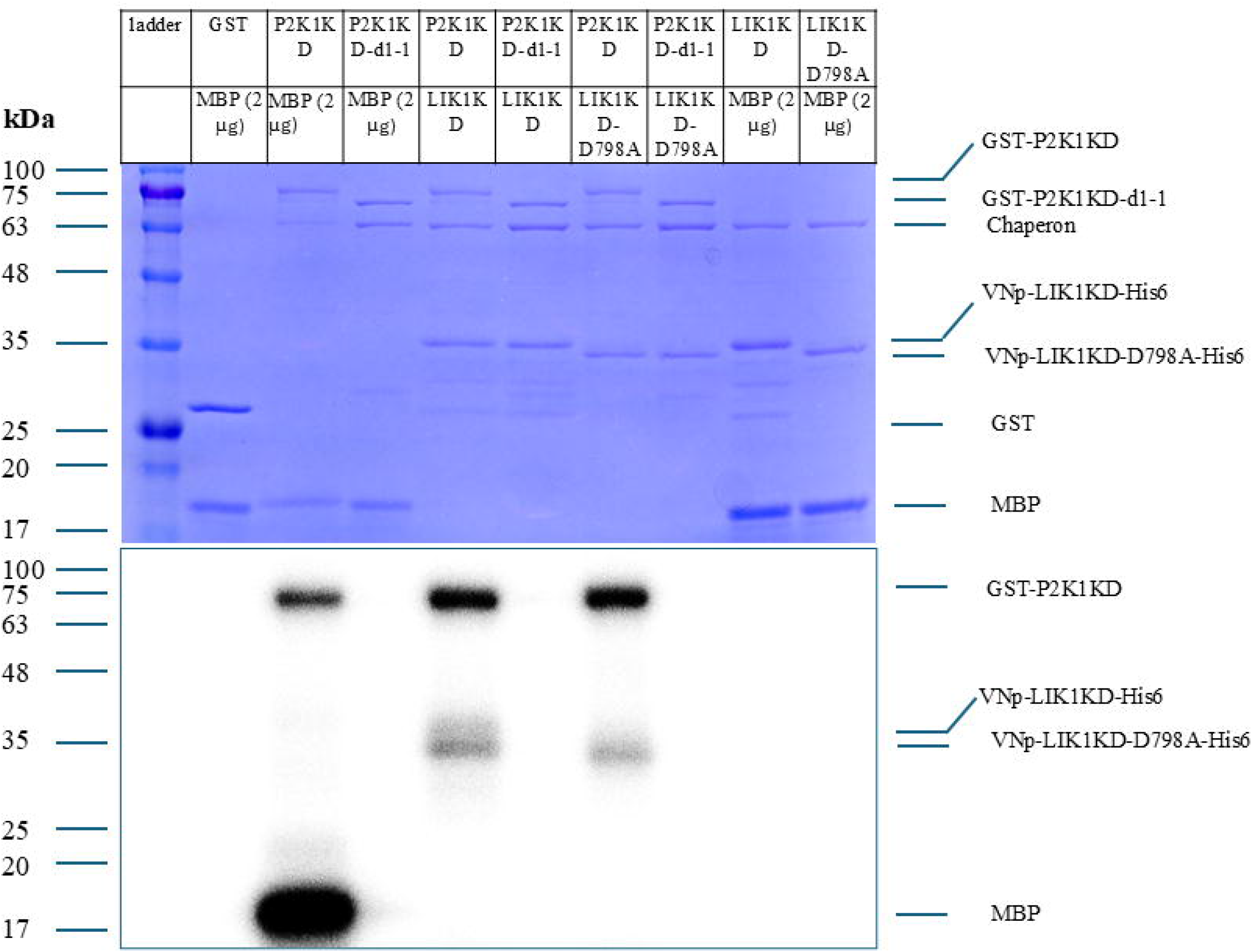
P2K1 can phosphorylate LIK1. Top: Coomassie brilliant blue staining to show the input proteins in the *in vitro* kinase assay in the presence of ^32^P labeled-ATP. Bottom: Radiograph to show the phosphorylation of the tested proteins. In this experiment, P2K1 kinase domain was fused to GST using the vector pGEX-5X-1 to form the GST-P2K1KD fusion protein, abbreviated as P2K1KD; P2K1KD mutant D525B (kinase dead) was fused to GST using the vector pGEX-5X-1 to form the GST-P2K1KD-d1-1, abbreviated as P2K1KD-d1-1; LIK1 kinase domain was fused to VNp and His6 using the vector pRSFDuet-1_VNp-His6 to form the VNp-LIK1KD-His6, abbreviated as LIK1KD; LIK1 mutant D798A (kinase dead) was fused to VNp and His6 using the vector pRSFDuet-1_VNp-His6 to form the VNp-LIK1KD-D798A-His6, abbreviated as LIK1KD-D798A. MBP (Myelin Basic Protein) was included as a positive kinase substrate. GST (Glutathione S-transferase) was included as a negative kinase substrate control. The experiment was repeated three times with similar results.

### 3.4. LIK1 is localized to the plasma membrane

LIK1 is predicted to localize to the plasma membrane, and our BiFC experiments supported this prediction by showing fluorescence at the cell periphery. To further confirm the subcellular localization of LIK1, we co-expressed a LIK1-GFP fusion construct with a plasma membrane marker (PM-mCherry) in *Nicotiana benthamiana* leaves. As shown in Fig. 6, the fluorescence signals of LIK1-GFP and PM-mCherry overlapped extensively, indicating LIK1’s localization at the plasma membrane. Furthermore, plasmolysis caused the LIK1-GFP signal to retract from the cell wall (Fig. S8), confirming its association with the plasma membrane rather than the cell wall. These results support that LIK1 is localized to the plasma membrane.

**Fig. 6.**
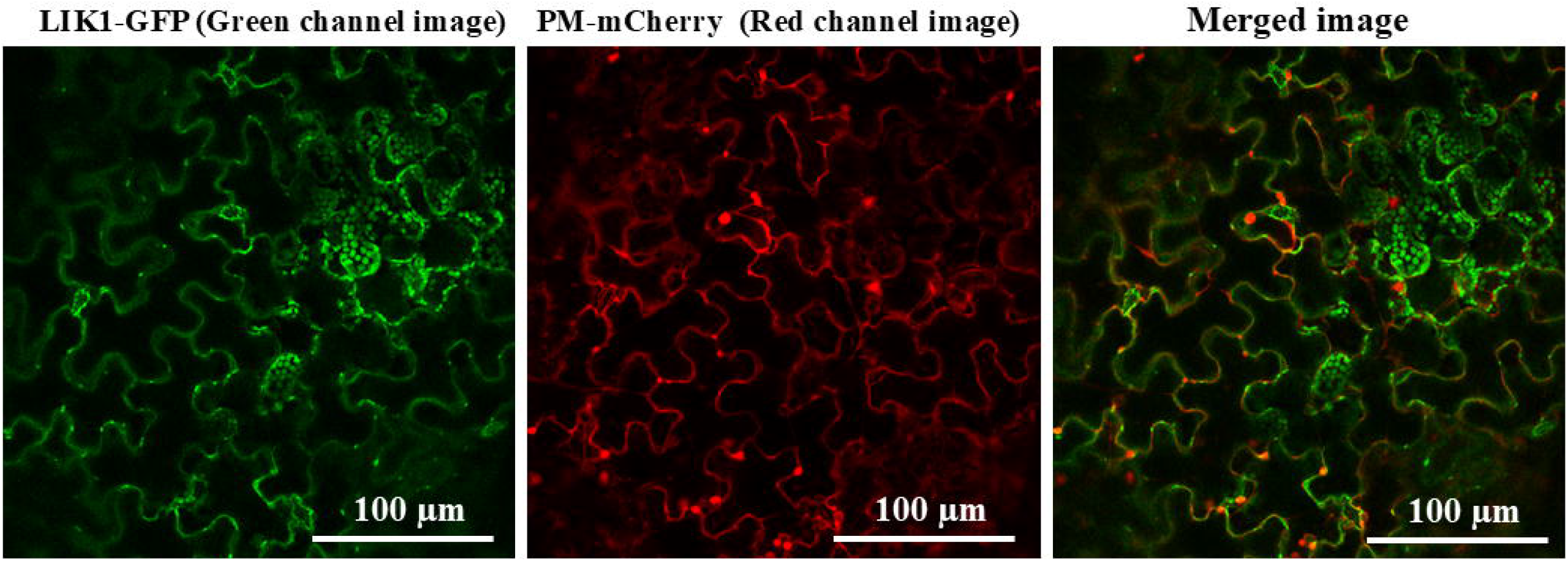
Localization of LIK1 to the plasma membrane. Tobacco (*Nicotiana benthamiana)* leaves were co-infiltrated with the following agrobacterial strain pairs containing: LIK1-GFP/PM-mCherry. The infiltrated leaves were observed ∼ three days later under a confocal microscope. Scale bars = 100 μm.

### 3.5. LIK1 is expressed ubiquitously and is not induced by ATP

To investigate the spatial expression pattern of *LIK1*, we generated transgenic Arabidopsis plants expressing a *LIK1* promoter-GUS reporter construct. Histochemical staining revealed that *LIK1* was ubiquitously expressed throughout the seedling, with relatively higher expression in leaves compared to roots (Fig. 7). No noticeable changes in GUS staining were observed in *LIK1* promoter–GUS transgenic plants following ATP treatment (Fig. S9). Consistently, quantitative RT-PCR analysis showed that *LIK1* transcript levels remained unchanged upon ATP treatment (Fig. S10). These results suggest that *LIK1* is constitutively expressed and not transcriptionally regulated by ATP.

**Fig. 7.**
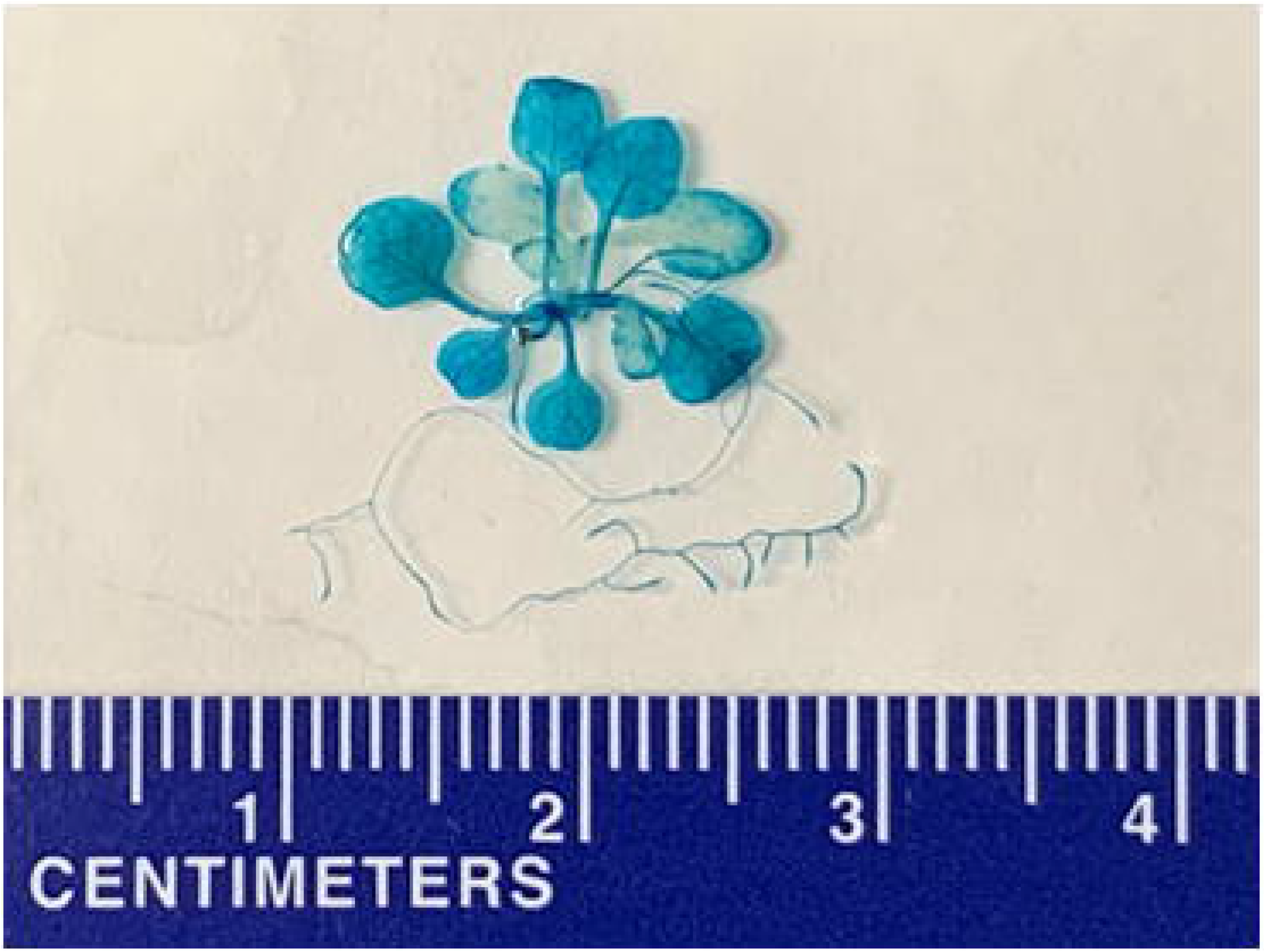
Histochemical analysis of GUS activity under the control of the *LIK1* gene promoter in transgenic plants. Ten-day-old seedlings transformed with the *LIK1* promoter-GUS construct were used in the analysis. A ruler was included in the picture as the scale bar: the distance between two neighboring lines is 1 millimeter.

## 4. Discussion

In this study, we demonstrated that mutations in the chitin receptor-interacting protein kinase gene *LIK1* significantly reduced the expression of eATP-responsive genes. We further showed that the LIK1 protein physically interacted with, and was phosphorylated by, the eATP receptor P2K1. These findings support our hypothesis that LIK1 functions in eATP signaling.

In our previous work, we reported that LIK1 interacted with the chitin receptor LysM RLK1/CERK1 to negatively regulate chitin signaling while positively modulating the JA signaling pathway (Le et al., 2014). These findings suggest that LIK1 may play a broader role in coordinating multiple signaling pathways.

eATP has been implicated in various developmental processes and in plant responses to both biotic and abiotic stresses (Cho et al., 2017; Roux and Steinebrunner, 2007; Yuan et al., 2025). Given these roles, it is not surprising that eATP signaling interacts with other pathways, particularly those involving plant hormones. Indeed, increasing evidence supports a significant overlap between the JA and eATP signaling pathways (Choi et al., 2014a; Tripathi et al., 2018; Tripathi and Tanaka, 2018; Ma et al., 2025).

As previously noted, LIK1 positively regulated the JA pathway (Le et al., 2014), which itself overlaps with eATP signaling. It is therefore reasonable to hypothesize that LIK1 may also serve as a positive regulator of eATP signaling. Indeed, our current data support this hypothesis, positioning LIK1 as a potential positive regulator of the eATP signaling pathway in *Arabidopsis thaliana*.

Nevertheless, the exact mechanism by which LIK1 influences eATP signaling remains to be elucidated. LIK1 is an LRR receptor-like kinase with a transmembrane domain, and our data confirmed its localization at the plasma membrane and its physical interaction with P2K1. Their physical interaction was also supported by the previous finding that LIK1 co-precipitated with P2K1 in transgenic *Arabidopsis* plants expressing P2K1-GFP (Sowders et al., 2024). Based on these observations, we propose that LIK1 and P2K1 may form part of a plasma membrane-localized receptor complex or nanodomain to regulate plant responses to eATP.

Receptor complexes at the plant cell membranes are increasingly recognized as critical hubs for integrating diverse signals that regulate growth, development, and stress responses. Their assembly and composition are dynamic, influenced by both internal and external cues (Burkart and Stahl, 2017). For instance, the FLS2-BAK1 complex mediates flagellin recognition to trigger immune responses (Chinchilla et al., 2007), whereas the BRI1-BAK1 complex perceives brassinosteroids to regulate growth (Li et al., 2002), and the LRK10L3-BAK1 complex senses eATP to mediate eATP-regulated seedling growth in Arabidopsis (Dong et al., 2025). Such complexes may include additional receptors and regulatory proteins, and their composition is likely shaped by stimuli to allow plants to respond appropriately (Burkart and Stahl, 2017; Bücherl et al., 2017; Hutten et al., 2017; Traeger et al., 2023; Wang et al., 2024).

Like BAK1, LIK1 may participate in distinct receptor complexes that perceive different signals and elicit appropriate downstream responses. Our previous and current data suggest that LIK1 is a component of the chitin receptor complex, where it negatively regulates chitin signaling, and also of the eATP receptor complex, where it positively regulates eATP signaling. Further studies are needed to characterize these receptor complexes in greater detail.

Beyond its interactions with LysM RLK1/CERK1 and P2K1, LIK1 has been reported to interact with 38 additional proteins, as documented in the Biological General Repository for Interaction Datasets (BioGRID) (Fig. S11). Notably, most of these interactors are leucine-rich repeat receptor kinases (Smakowska-Luzan et al., 2018), implying that LIK1 may engage in additional receptor complexes and participate in other signaling pathways.

In summary, our findings, together with existing data, support the role of LIK1 as a novel regulator of eATP signaling, in addition to its previously established function in chitin signaling. This suggests that LIK1 may serve as a versatile signaling component within different receptor complexes. By interacting with different proteins in these complexes, LIK1 may help coordinate plant responses to a broad spectrum of internal and external cues, including chitin and eATP.

## Supporting information

Table S2. Primers used in this study

Fig. S11. The interactions of LIK1 with different proteins

Fig. S1. The LIK1 T-DNA insertion mutant (SALK_030855, lik1-1)

Fig. S2. qRT-PCR analysis of ATP-responsive genes in lik1-1

Fig. S3. DNA sequence chromatogram to show the 1-bp insertion in the LIK1 gene in the edited line lik1-2

Fig. S4. One base insertion in lik1-2 led to an early stop codon in the LIK1 CDS

Fig. S5. DNA sequence chromatogram to show the 1-bp deletion in the LIK1 edited line lik1-3

Fig. S6. One base deletion in lik1-3 led to an early stop codon in the LIK1 CDS

Fig. S7. LIK1-P2K1 interaction in the bimolecular fluorescence complementation (BiFC) assay

Fig. S8. Plasmolysis to show the separation of the LIK1-GFP signal from the cell wall

Fig. S10. The expression of LIK1 in response to ATP

Fig. S9. Histochemical analysis of GUS activity under the control of the LIK1 gene promoter in transgenic plants in response to ATP treatment

Table S1. The peptides from LIK1 showed differential abundances between the ATP and mock treatments in the limited proteolysis (LiP) proteomics analys

## Abbreviations

DAMP: damage-associated molecular pattern
eATP: extracellular ATP
JA: jasmonic acid
LIK1: LysM RLK1-interacting kinase 1
P2K1: purinergic receptor kinase 1.

## CRediT authorship contribution statement

**Jinrong Wan:** data acquisition, analysis, and writing. **Mengran Yang:** data acquisition, manuscript review and editing. **Jae Song:** data acquisition, manuscript review and editing. **Chunhui Xu**: data analysis, manuscript review and editing. **Sung-Hwan Cho:** data acquisition, manuscript review and editing. **Mowei Zhou:** data acquisition and analysis. **Ljiljana Pasa-Tolic:** data acquisition, analysis and supervision. **Bing Yang**: gene editing instructions. **Dong Xu**: data analysis, manuscript review and editing. **Gary Stacey**: funding acquisition, supervision, and manuscript revision.

## Supporting information

The following is the Supplementary material related to this article.

Table S1. The peptides from LIK1 showed differential abundances between the ATP and mock treatments in the limited proteolysis (LiP) proteomics analysis.

Table S2. Primers used in this study.

Fig. S1. The *LIK1* T-DNA insertion mutant (SALK_030855, *lik1-1*). (A) Diagrammatic representation of the T-DNA insertion in the *LIK1* gene. The diagram was not drawn to scale. (B) RT-PCR analysis of the homozygous *lik1-1* mutant. M: 1-Kb DNA ladder; WT: wild-type Arabidopsis plants; *lik1-1*: homozygous SALK_030855 mutant; *SAND*: the reference gene *SAND* (AT2G28390) used as the control.

Fig. S2. qRT-PCR analysis of ATP-responsive genes in *lik1-1*. Top: *CPK28*. Bottom: *WRKY40*. One-week-old seedlings were treated with 200 µM ATP or mock control (MES buffer) for 30 minutes. The relative expression (fold change) of a target gene was obtained by comparing the ATP-treated samples to mock controls, following normalization to the reference gene *SAND*. Data represented the mean ± SE (standard error) of four biological replicates. Statistical significance (between the WT and *lik1-1*) was determined using a Student’s t-test: **, *P* value <0.01.

Fig. S3. DNA sequence chromatogram to show the 1-bp insertion in the *LIK1* gene in the edited line *lik1-2*. The arrow indicated the 1-bp insertion. The gRNA targeting sequence and the PAM (Protospacer Adjacent Motif, in bold and green) were included to show the position of the insertion in this edited line.

Fig. S4. One base insertion in *lik1-2* led to an early stop codon in the *LIK1* CDS. The inserted base, A, was highlighted in bold and red. The original start and stop codons were highlighted in bold. The new stop codon TAA was highlighted in bold and red.

Fig. S5. DNA sequence chromatogram to show the 1-bp deletion in the *LIK1* edited line *lik1-3*. The arrow indicated the position of the 1-bp deletion. The gRNA targeting sequence and the PAM (Protospacer Adjacent Motif, in bold and green) were included to show the position of the deletion in this edited line.

Fig. S6. One base deletion in *lik1-3* led to an early stop codon in the *LIK1* CDS. The deleted base, A, was highlighted in bold and grey. The original start and stop codons were highlighted in bold. The new stop codon TAG was highlighted in bold and red.

Fig. S7. LIK1-P2K1 interaction in the bimolecular fluorescence complementation (BiFC) assay. Tobacco (*Nicotiana benthamiana)* leaves were co-infiltrated with the following agrobacterial strain pairs containing: LIK1-NYFP/P2K1-CYFP, LIK1-NYFP/CYFP, NYFP/P2K1-CYFP, and NYFP/CYFP, as indicated in the figure. The infiltrated leaves were observed ∼ three days later under a confocal microscope. Scale bars = 100 μm.

Fig. S8. Plasmolysis to show the separation of the LIK1-GFP signal from the cell wall. Tobacco (*Nicotiana benthamiana)* leaves were infiltrated with the agrobacterial strain containing: LIK1-GFP. Three days later, the infiltrated leaves were collected and treated with 3 N NaCl for ∼15 minutes before observation under a confocal microscope. Scale bars = 100 μm.

Fig. S9. Histochemical analysis of GUS activity under the control of the *LIK1* gene promoter in transgenic plants in response to ATP treatment. Ten-day-old seedlings transformed with the *LIK1* promoter-GUS construct were treated with ATP (200 µM) or mock (MES buffer) for 30 minutes. A ruler was included in each picture as the scale bar: the distance between two neighboring lines is 1 millimeter.

Fig. S10. The expression of *LIK1* in response to ATP. One-week-old Wild-type seedlings were treated with 200 µM ATP or mock control (MES buffer) for 30 minutes. Data represented the mean ± SE (standard error, n=4) after normalization to the reference gene *SAND*. Statistical significance (between the ATP-treated and mock samples) was determined using a Student’s t-test: no statistical significance was found.

Fig. S11. The interactions of LIK1 with different proteins. The data were from: https://thebiogrid.org/6047/summary/arabidopsis-thaliana/at3g14840.html

## Data availability

Data will be made available on request.

## Acknowledgements

Not applicable.

## References

Bücherl CA, Jarsch IK, Schudoma C, Segonzac C, Mbengue M, Robatzek S, MacLean D, Ott T, Zipfel C. Plant immune and growth receptors share common signalling components but localise to distinct plasma membrane nanodomains. Elife. 2017 Mar 6;6:e25114. doi: 10.7554/eLife.25114. PMID: 28262094; PMCID: PMC5383397.

Burkart RC, Stahl Y. Dynamic complexity: plant receptor complexes at the plasma membrane. Curr Opin Plant Biol. 2017 Dec;40:15–21. doi: 10.1016/j.pbi.2017.06.016. Epub 2017 Jul 14. PMID: 28715768.

Cappelletti V, Hauser T, Piazza I, Pepelnjak M, Malinovska L, Fuhrer T, Li Y, Dörig C, Boersema P, Gillet L, Grossbach J, Dugourd A, Saez-Rodriguez J, Beyer A, Zamboni N, Caflisch A, de Souza N, Picotti P. Dynamic 3D proteomes reveal protein functional alterations at high resolution in situ. Cell. 2021 Jan 21;184(2):545–559.e22. doi: 10.1016/j.cell.2020.12.021. Epub 2020 Dec 23. PMID: 33357446; PMCID: PMC7836100.

Chen D, Cao Y, Li H, Kim D, Ahsan N, Thelen J, Stacey G. Extracellular ATP elicits DORN1-mediated RBOHD phosphorylation to regulate stomatal aperture. Nat Commun. 2017 Dec 22;8(1):2265. doi: 10.1038/s41467-017-02340-3. PMID: 29273780; PMCID: PMC5741621.

Chen H, Zou Y, Shang Y, Lin H, Wang Y, Cai R, Tang X, Zhou JM. Firefly luciferase complementation imaging assay for protein-protein interactions in plants. Plant Physiol. 2008 Feb;146(2):368–76. doi: 10.1104/pp.107.111740. Epub 2007 Dec 7. PMID: 18065554; PMCID: PMC2245818.

Chinchilla D, Zipfel C, Robatzek S, Kemmerling B, Nürnberger T, Jones JD, Felix G, Boller T. A flagellin-induced complex of the receptor FLS2 and BAK1 initiates plant defence. Nature. 2007 Jul 26;448(7152):497–500. doi: 10.1038/nature05999. Epub 2007 Jul 11. PMID: 17625569.

Cho SH, Nguyen CT, Choi J, Stacey G. Molecular Mechanism of Plant Recognition of Extracellular ATP. Adv Exp Med Biol. 2017;1051:233–253. doi: 10.1007/5584_2017_110. PMID: 29064066.

Cho SH, Tóth K, Kim D, Vo PH, Lin CH, Handakumbura PP, Ubach AR, Evans S, Paša-Tolić L, Stacey G. Activation of the plant mevalonate pathway by extracellular ATP. Nat Commun. 2022 Jan 21;13(1):450. doi: 10.1038/s41467-022-28150-w. PMID: 35064110; PMCID: PMC8783019.

Choi J, Tanaka K, Cao Y, Qi Y, Qiu J, Liang Y, Lee SY, Stacey G. Identification of a plant receptor for extracellular ATP. Science. 2014a Jan 17;343(6168):290–4. doi: 10.1126/science.343.6168.290. Erratum in: Science. 2014 Feb 14;343(6172):730. PMID: 24436418.

Choi J, Tanaka K, Liang Y, Cao Y, Lee SY, Stacey G. Extracellular ATP, a danger signal, is recognized by DORN1 in Arabidopsis. Biochem J. 2014b Nov 1;463(3):429–37. doi: 10.1042/BJ20140666. PMID: 25301072.

Clough SJ, Bent AF. Floral dip: a simplified method for Agrobacterium-mediated transformation of Arabidopsis thaliana. Plant J. 1998 Dec;16(6):735–43. doi: 10.1046/j.1365-313x.1998.00343.x. PMID: 10069079.

Curtis MD, Grossniklaus U. A gateway cloning vector set for high-throughput functional analysis of genes in planta. Plant Physiol. 2003 Oct;133(2):462–9. doi: 10.1104/pp.103.027979. PMID: 14555774; PMCID: PMC523872.

Dong X, Xu J, Fu Y, Wang T, Yin H, Li Y, Yu L, Zhu R, Kang E, Shang Z. LRK10L3 and BAK1 are collaboratively involved in extracellular ATP-regulated seedling growth of Arabidopsis thaliana. New Phytol. 2025 Jun 29. doi: 10.1111/nph.70352. Epub ahead of print. PMID: 40583302.

Eastwood TA, Baker K, Streather BR, Allen N, Wang L, Botchway SW, Brown IR, Hiscock JR, Lennon C, Mulvihill DP. High-yield vesicle-packaged recombinant protein production from E. coli. Cell Rep Methods. 2023 Feb 2;3(2):100396. doi: 10.1016/j.crmeth.2023.100396. PMID: 36936078; PMCID: PMC10014274.

Hooper CM, Castleden IR, Tanz SK, Aryamanesh N, Millar AH. SUBA4: the interactive data analysis centre for Arabidopsis subcellular protein locations. Nucleic Acids Res. 2017 Jan 4;45(D1):D1064–D1074. doi: 10.1093/nar/gkw1041. Epub 2016 Nov 28. PMID:27899614; PMCID: PMC5210537.

Hooper CM, Tanz SK, Castleden IR, Vacher MA, Small ID, Millar AH. SUBAcon: a consensus algorithm for unifying the subcellular localization data of the Arabidopsis proteome. Bioinformatics. 2014 Dec 1;30(23):3356–64. doi:10.1093/bioinformatics/btu550. Epub 2014 Aug 22. PMID: 25150248.

Hutten SJ, Hamers DS, Aan den Toorn M, van Esse W, Nolles A, Bücherl CA, de Vries SC, Hohlbein J, Borst JW. Visualization of BRI1 and SERK3/BAK1 Nanoclusters in Arabidopsis Roots. PLoS One. 2017 Jan 23;12(1):e0169905. doi: 10.1371/journal.pone.0169905. PMID: 28114413; PMCID: PMC5256950.

Jefferson RA, Kavanagh TA, Bevan MW. GUS fusions: beta-glucuronidase as a sensitive and versatile gene fusion marker in higher plants. EMBO J. 1987 Dec 20;6(13):3901–7. doi: 10.1002/j.1460-2075.1987.tb02730.x. PMID: 3327686; PMCID: PMC553867.

Jewell JB, Carlton A, Bartley LE, Tanaka K. Jasmonate primes plant responses to extracellular ATP through purinoceptor P2K1. bioRxiv 2024.11.07.622526; doi:10.1101/2024.11.07.622526.

Jorge GL, Kim D, Xu C, Cho SH, Su L, Xu D, Bartley LE, Stacey G, Thelen JJ. Unveiling orphan receptor-like kinases in plants: novel client discovery using high-confidence library predictions in the Kinase-Client (KiC) assay. Front Plant Sci. 2024 Apr 3;15:1372361. doi: 10.3389/fpls.2024.1372361. PMID: 38633461; PMCID: PMC11021772.

Kim D, Jorge GL, Xu C, Su L, Cho SH, Ahsan N, Chen D, Zhou L, Gritsenko MA, Zhou M, Wan J, Pasa-Tolic L, Xu D, Bartley LE, Thelen JJ, Stacey G. Identifying Receptor Kinase Substrates Using an 8000 Peptide Kinase Client Library Enriched for Conserved Phosphorylation Sites. Mol Cell Proteomics. 2025 Mar;24(3):100926. doi: 10.1016/j.mcpro.2025.100926. Epub 2025 Feb 7. PMID: 39923935; PMCID: PMC11952801.

Kimberlin AN, Holtsclaw RE, Zhang T, Mulaudzi T, Koo AJ. On the initiation of jasmonate biosynthesis in wounded leaves. Plant Physiol. 2022 Aug 1;189(4):1925–1942. doi: 10.1093/plphys/kiac163. PMID: 35404431; PMCID: PMC9342990.

Koo AJ, Howe GA. The wound hormone jasmonate. Phytochemistry. 2009 Sep;70(13-14):1571–80. doi: 10.1016/j.phytochem.2009.07.018. Epub 2009 Aug 18. PMID:19695649; PMCID: PMC2784233.

Le MH, Cao Y, Zhang XC, Stacey G. LIK1, a CERK1-interacting kinase, regulates plant immune responses in Arabidopsis. PLoS One. 2014 Jul 18;9(7):e102245. doi: 10.1371/journal.pone.0102245. PMID: 25036661; PMCID: PMC4103824.

Li J, Wen J, Lease KA, Doke JT, Tax FE, Walker JC. BAK1, an Arabidopsis LRR receptor-like protein kinase, interacts with BRI1 and modulates brassinosteroid signaling. Cell. 2002 Jul 26;110(2):213–22. doi: 10.1016/s0092-8674(02)00812-7. PMID: 12150929.

Li X. Infiltration of Nicotiana benthamiana protocol for transient expression via Agrobacterium. Bio-protocol. 2011;1(e95).

Livak KJ, Schmittgen TD. Analysis of relative gene expression data using real-time quantitative PCR and the 2(-Delta Delta C(T)) Method. Methods. 2001 Dec;25(4):402–8. doi: 10.1006/meth.2001.1262. PMID: 11846609.

Ma X, Hasan MS, Anjam MS, Mahmud S, Bhattacharyya S, Vothknecht UC, Mendy B, Grundler FMW, Marhavý P. Ca2+ waves and ethylene/JA crosstalk orchestrate wound responses in Arabidopsis roots. EMBO Rep. 2025 May 19. doi: 10.1038/s44319-025-00471-z. Epub ahead of print. PMID: 40389757.

Miya A, Albert P, Shinya T, Desaki Y, Ichimura K, Shirasu K, Narusaka Y, Kawakami N, Kaku H, Shibuya N. CERK1, a LysM receptor kinase, is essential for chitin elicitor signaling in Arabidopsis. Proc Natl Acad Sci U S A. 2007 Dec 4;104(49):19613–8. doi: 10.1073/pnas.0705147104. Epub 2007 Nov 27. PMID: 18042724; PMCID: PMC2148337.

Myers RJ, Fichman Y, Stacey G, Mittler R. Extracellular ATP plays an important role in systemic wound response activation. Plant Physiol. 2022 Jun 27;189(3):1314–1325. doi: 10.1093/plphys/kiac148. PMID: 35348752; PMCID: PMC9237675.

Pham AQ, Cho SH, Nguyen CT, Stacey G. Arabidopsis Lectin Receptor Kinase P2K2 Is a Second Plant Receptor for Extracellular ATP and Contributes to Innate Immunity. Plant Physiol. 2020 Jul;183(3):1364–1375. doi: 10.1104/pp.19.01265. Epub 2020 Apr 28. PMID: 32345768; PMCID: PMC7333714.

Roux SJ, Steinebrunner I. Extracellular ATP: an unexpected role as a signaler in plants. Trends Plant Sci. 2007 Nov;12(11):522–527. doi: 10.1016/j.tplants.2007.09.003. Epub 2007 Oct 24. PMID: 17928260.

Silhavy D, Molnár A, Lucioli A, Szittya G, Hornyik C, Tavazza M, Burgyán J. A viral protein suppresses RNA silencing and binds silencing-generated, 21-to 25-nucleotide double-stranded RNAs. EMBO J. 2002 Jun 17;21(12):3070–80. doi: 10.1093/emboj/cdf312. PMID: 12065420; PMCID: PMC125389.

Smakowska-Luzan E, Mott GA, Parys K, Stegmann M, Howton TC, Layeghifard M, Neuhold J, Lehner A, Kong J, Grünwald K, Weinberger N, Satbhai SB, Mayer D, Busch W, Madalinski M, Stolt-Bergner P, Provart NJ, Mukhtar MS, Zipfel C, Desveaux D, Guttman DS, Belkhadir Y. An extracellular network of Arabidopsis leucine-rich repeat receptor kinases. Nature. 2018 Jan 18;553(7688):342–346. doi: 10.1038/nature25184. Epub 2018 Jan 10. Erratum in: Nature. 2018 Sep;561(7722):E8. doi: 10.1038/s41586-018-0268-y. PMID: 29320478; PMCID: PMC6485605.

Sowders JM, Jewell JB, Tanaka K. CPK28 is a modulator of purinergic signaling in plant growth and defense. Plant J. 2024 May;118(4):1086–1101. doi: 10.1111/tpj.16656. Epub 2024 Feb 3. PMID: 38308597; PMCID: PMC11096078.

Tanaka K, Choi J, Cao Y, Stacey G. Extracellular ATP acts as a damage-associated molecular pattern (DAMP) signal in plants. Front Plant Sci. 2014 Sep 3;5:446. doi: 10.3389/fpls.2014.00446. PMID: 25232361; PMCID: PMC4153020.

Tanaka K, Heil M. Damage-Associated Molecular Patterns (DAMPs) in Plant Innate Immunity: Applying the Danger Model and Evolutionary Perspectives. Annu Rev Phytopathol. 2021 Aug 25;59:53–75. doi: 10.1146/annurev-phyto-082718-100146. Epub 2021 Apr 26. PMID: 33900789.

Tripathi D, Tanaka K. A crosstalk between extracellular ATP and jasmonate signaling pathways for plant defense. Plant Signal Behav. 2018;13(5):e1432229. doi: 10.1080/15592324.2018.1432229. Epub 2018 Feb 20.

Tripathi D, Zhang T, Koo AJ, Stacey G, Tanaka K. Extracellular ATP Acts on Jasmonate Signaling to Reinforce Plant Defense. Plant Physiol. 2018 Jan;176(1):511–523. doi: 10.1104/pp.17.01477. Epub 2017 Nov 27. PMID: 29180381; PMCID: PMC6108377.

Wan J, Zhang XC, Neece D, Ramonell KM, Clough S, Kim SY, Stacey MG, Stacey G. A LysM receptor-like kinase plays a critical role in chitin signaling and fungal resistance in Arabidopsis. Plant Cell. 2008 Feb;20(2):471–81. doi: 10.1105/tpc.107.056754. Epub 2008 Feb 8. PMID: 18263776; PMCID: PMC2276435.

Wang B, Zhou Z, Zhou JM, Li J. Myosin XI-mediated BIK1 recruitment to nanodomains facilitates FLS2-BIK1 complex formation during innate immunity in Arabidopsis. Proc Natl Acad Sci U S A. 2024 Jun 18;121(25):e2312415121. doi: 10.1073/pnas.2312415121. Epub 2024 Jun 14. PMID: 38875149; PMCID: PMC11194512.

Yuan DP, Kim D, Xuan YH (2025), Extracellular ATP: an emerging multifaceted regulator of plant fitness. Plant Biotechnol. J. 2025 Feb 12; 23(5):1771–1782. 10.1111/pbi.70006

Zhou L, Zhou M, Gritsenko MA, Stacey G. Selective Enrichment Coupled with Proteomics to Identify S-Acylated Plasma Membrane Proteins in Arabidopsis. Curr Protoc Plant Biol. 2020 Dec;5(4):e20119. doi: 10.1002/cppb.20119. PMID: 32976704; PMCID: PMC9206763.

