## Supplementary figures and images for "The chitin receptor-interacting protein LIK1 regulates extracellular ATP signaling via interaction with P2K1 in *Arabidopsis thaliana*"

### Fig. S1. The LIK1 T-DNA insertion mutant (SALK_030855, lik1-1)

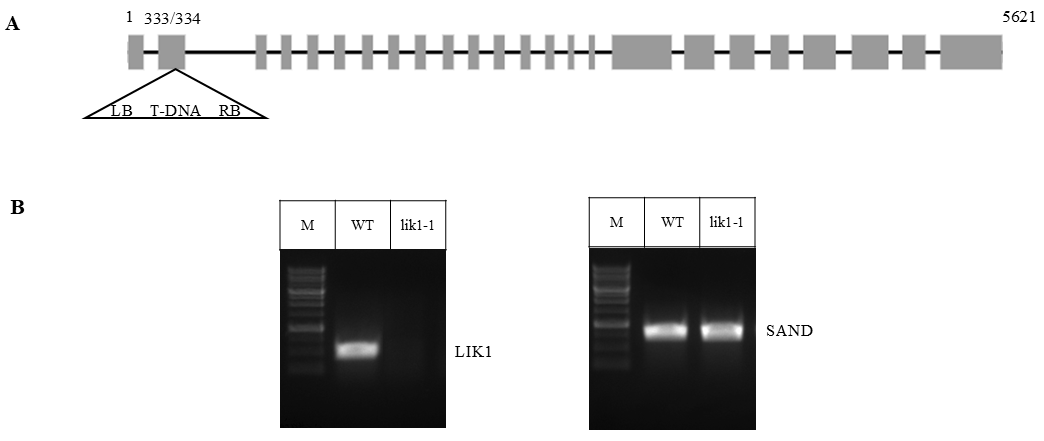

### Fig. S2. qRT-PCR analysis of ATP-responsive genes in lik1-1

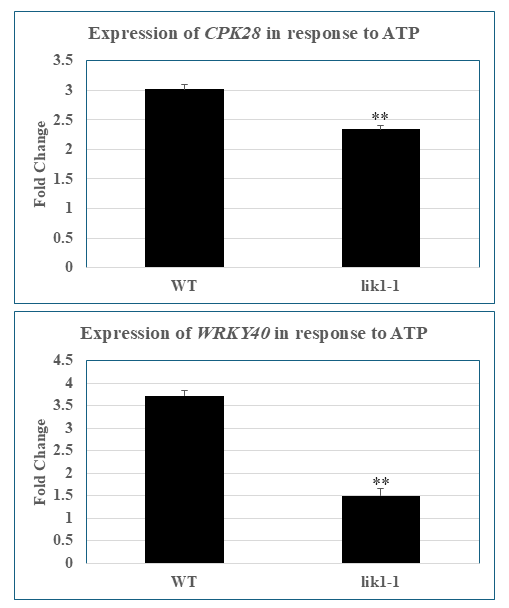

### Fig. S3. DNA sequence chromatogram to show the 1-bp insertion in the LIK1 gene in the edited line lik1-2

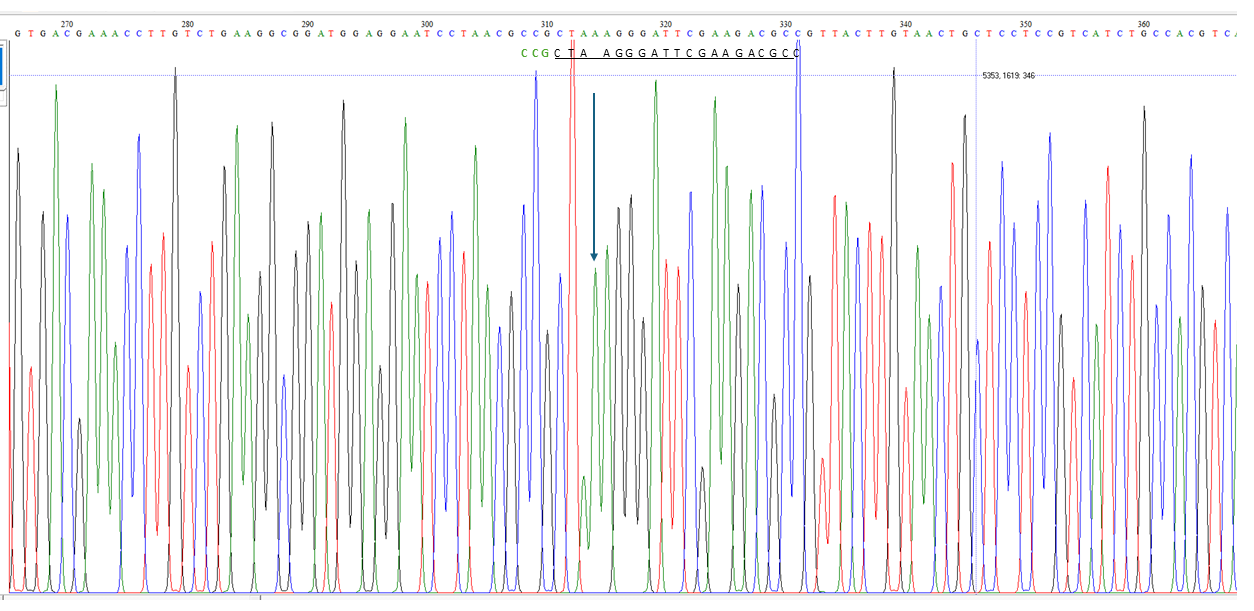

### Fig. S4. One base insertion in lik1-2 led to an early stop codon in the LIK1 CDS

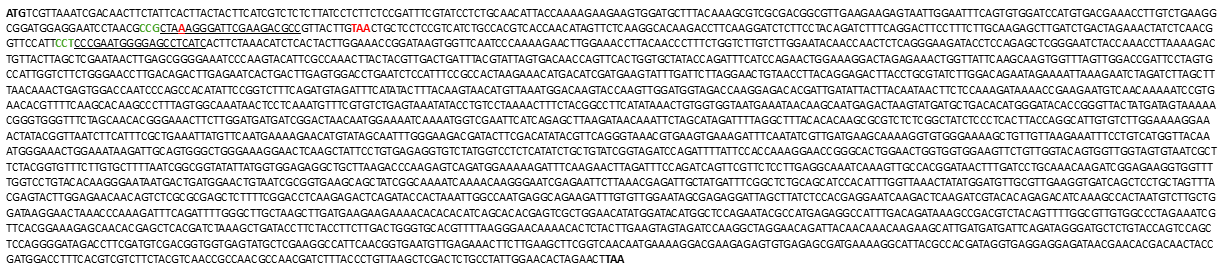

### Fig. S5. DNA sequence chromatogram to show the 1-bp deletion in the LIK1 edited line lik1-3

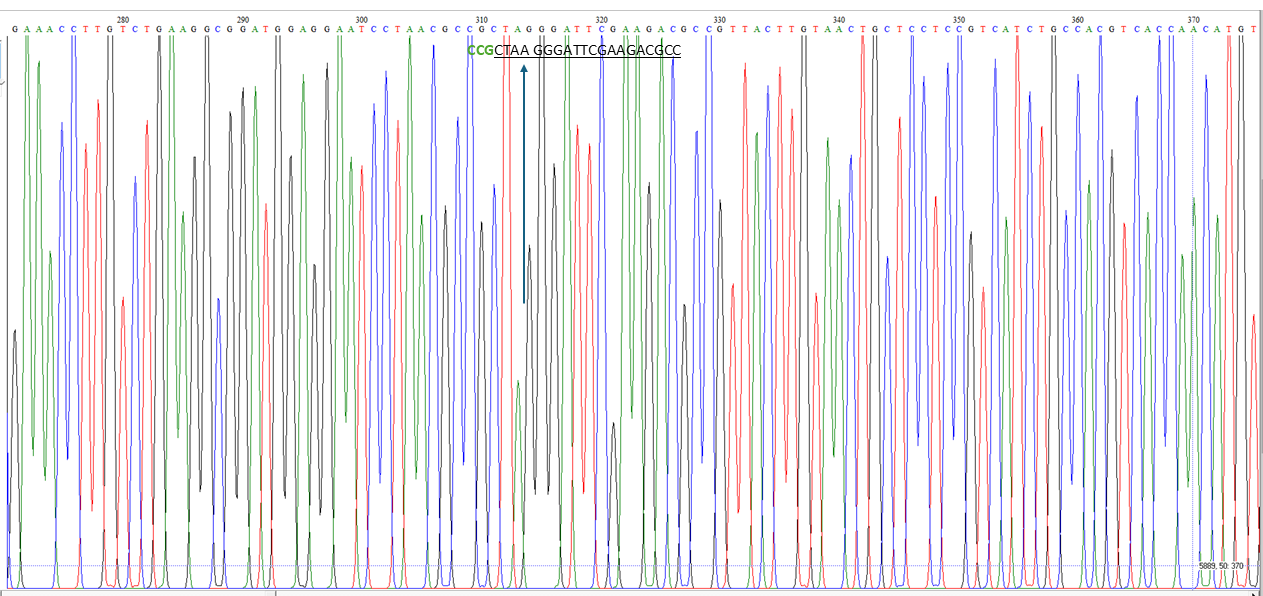

### Fig. S6. One base deletion in lik1-3 led to an early stop codon in the LIK1 CDS

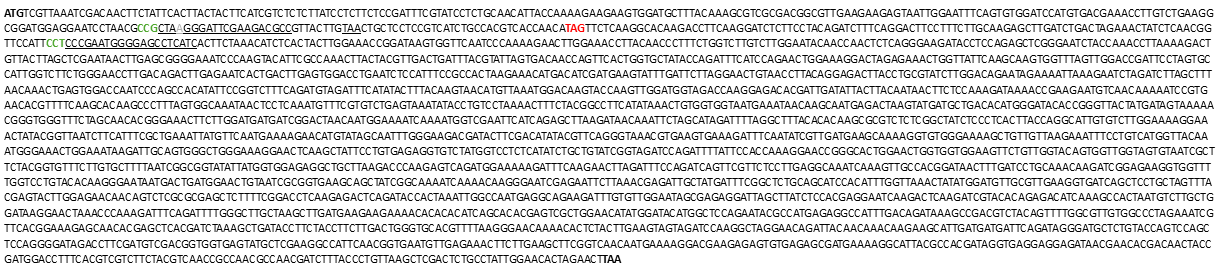

### Fig. S7. LIK1-P2K1 interaction in the bimolecular fluorescence complementation (BiFC) assay

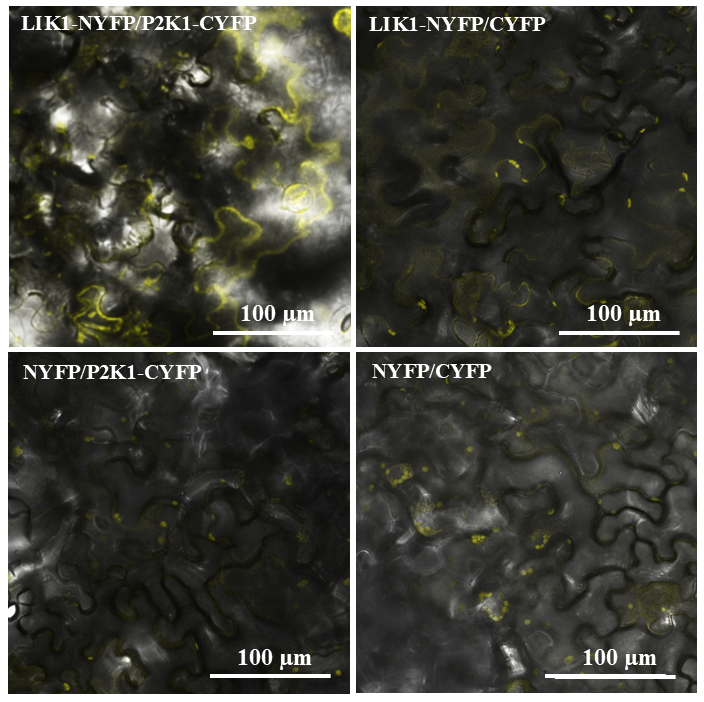

### Fig. S8. Plasmolysis to show the separation of the LIK1-GFP signal from the cell wall

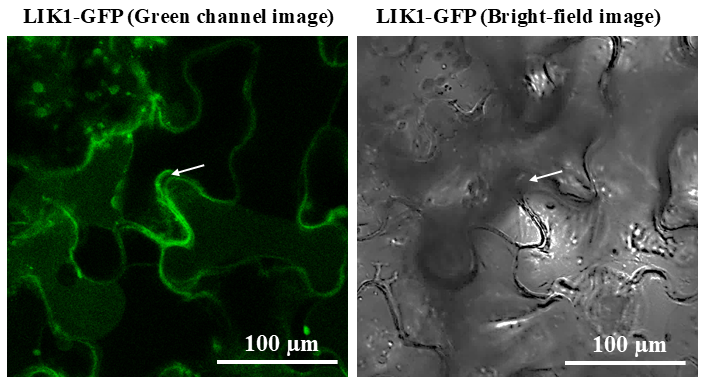

### Fig. S9. Histochemical analysis of GUS activity under the control of the LIK1 gene promoter in transgenic plants in response to ATP treatment

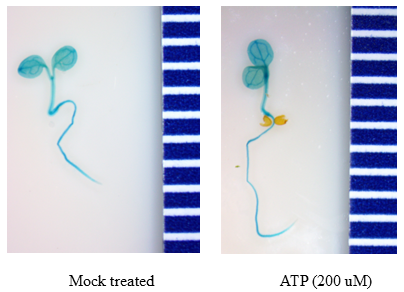

### Fig. S10. The expression of LIK1 in response to ATP

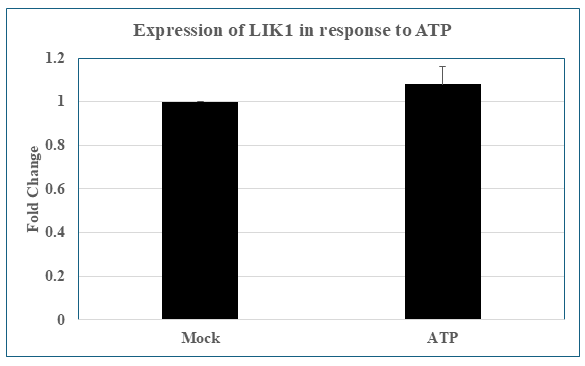

### Fig. S11. The interactions of LIK1 with different proteins

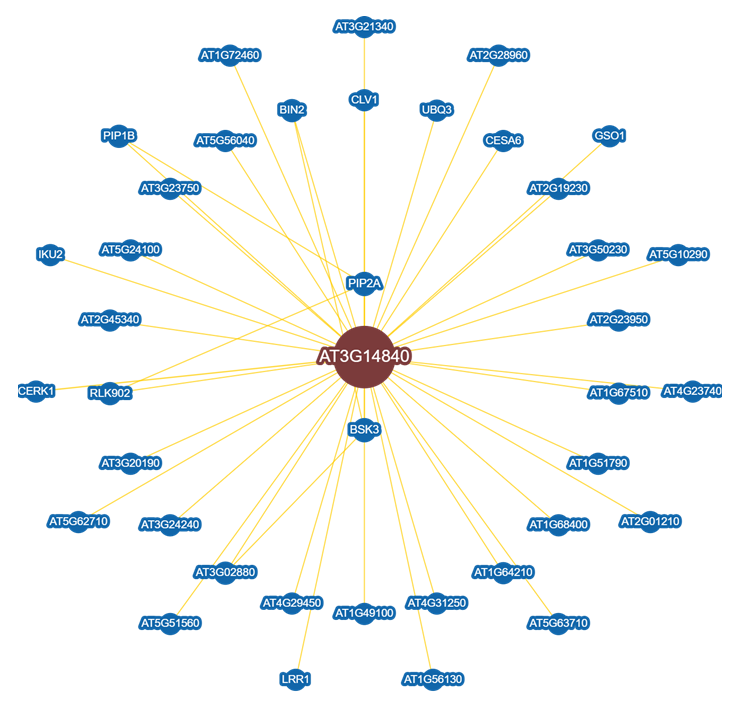
